# Transcriptional profiling of zebrafish identifies host factors controlling susceptibility to *Shigella flexneri*

**DOI:** 10.1101/2022.10.03.510593

**Authors:** Vincenzo Torraca, Richard J. White, Ian M. Sealy, Maria Mazon-Moya, Gina Duggan, Alexandra Willis, Elisabeth M. Busch-Nentwich, Serge Mostowy

## Abstract

*Shigella flexneri* is a human adapted pathovar of *Escherichia coli* that can invade the intestinal epithelium, causing inflammation and bacillary dysentery. Although an important human pathogen, the host response to *S. flexneri* is poorly understood. Zebrafish larvae, highly innovative for genomics, transcriptomics and genetic tractability, represent a valuable animal model to study human infections *in vivo*. Here we use a *S. flexneri*-zebrafish infection model to generate mRNA expression profiles of host response to *S. flexneri* infection at the whole animal level. The signature of early *S. flexneri* infection (detected at 6 hours post-infection) is dominated by immune response-related processes. Consistent with its clearance from the host, the signature of late *S. flexneri* infection (detected at 24 hours post-infection) is significantly changed, where only a small set of immune-related genes remain differentially expressed, including *gpr84* which encodes a putative G-protein coupled receptor. Using mutant zebrafish lines generated by ENU, CRISPR mutagenesis and the F0 CRISPR knockout method, we show that *gpr84*-deficient larvae are more susceptible to *S. flexneri* infection. Together, these results highlight the power of zebrafish to model infection by bacterial pathogens and provide a community resource to investigate host response to *S. flexneri* infection.

## INTRODUCTION

The zebrafish has emerged as an important animal model to study human infection (Torraca *et al*., 2018; Gomes and Mostowy, 2020; Stream and Madigan, 2022). Zebrafish larvae are genetically tractable, optically accessible and present a fully functional innate immune system with macrophages and neutrophils that mimic their mammalian counterparts. A wide variety of pathogenic bacteria have been investigated using zebrafish models, providing unprecedented resolution of the cellular response to infection *in vivo*.

*Shigella flexneri* is an important human pathogen and causative agent of bacillary dysentery. It is highly infectious and its prevalence is highest in tropical/subtropical regions of the world where access to safe drinking water is limited. Mice are naturally resistant to *S. flexneri* infection (Schnupf and Sansonetti, 2019), although new work has shown that NAIP–NLRC4-deficient mice are susceptible to oral *S. flexneri* infection and recapitulate clinical features of shigellosis (Mitchell *et al*., 2020). A *S. flexneri*-zebrafish infection model has been established, showing that both macrophages and neutrophils are involved in the protective immune response, and providing mechanistic insights into interactions between *S. flexneri* and phagocytes (Mostowy *et al*., 2013; Duggan and Mostowy, 2018; Torraca *et al*., 2018). The zebrafish infection model for *S. flexneri* has been shown to successfully recapitulate key features of shigellosis including inflammation and macrophage cell death (Mostowy *et al*., 2013; Mazon-Moya *et al*., 2017), and has been valuable in highlighting key roles of virulence factors (e.g. T3SS, O-antigen) (Mostowy *et al*., 2013; Mazon-Moya *et al*., 2017; Torraca *et al*., 2019) and cell-autonomous immunity (e.g. autophagy, the septin cytoskeleton) (Mostowy *et al*., 2013; Mazon-Moya *et al*., 2017) in host-pathogen interactions.

Here, using our *S. flexneri*-zebrafish infection model, we generate a high-resolution mRNA expression time course of host response to *S. flexneri* infection at the whole animal level. We discover a rapid, but largely transient, activation of immune-related processes in response to sublethal *S. flexneri* infection. Strikingly, only a discrete set of immune genes (including a gene encoding the putative G-protein coupled receptor Gpr84) remain differentially expressed at 24 hours post infection (hpi), suggesting an important role for these markers in host survival. Consistent with this, we show that *gpr84* mutation leads to increased susceptibility to *S. flexneri* infection. Our transcriptional profiles obtained using zebrafish provide a community resource to investigate host response to *S. flexneri* infection.

## RESULTS

### Whole animal RNA-seq profiling of *S. flexneri* infected larvae

To obtain the gene expression response of zebrafish larvae to *S. flexneri* infection we performed caudal vein injections at 2 days post fertilisation (dpf). We chose infection doses (1000 CFU) that are non-lethal by 24 hours post infection (hpi) to avoid eliciting non-specific transcriptional responses due to systemic stress. Consistent with this, *S. flexneri* burden decreased over time (**Fig. 1A**) and zebrafish larvae survived infections for the entire observation period of 120 hours (**Fig. 1B**). We collected four pools of five larvae, each from uninfected, mock-infected with PBS and infected groups at 6 hpi and 24 hpi, extracted RNA and produced sequencing libraries for RNA-seq **(Fig. 1C)**.

**Figure 1.**
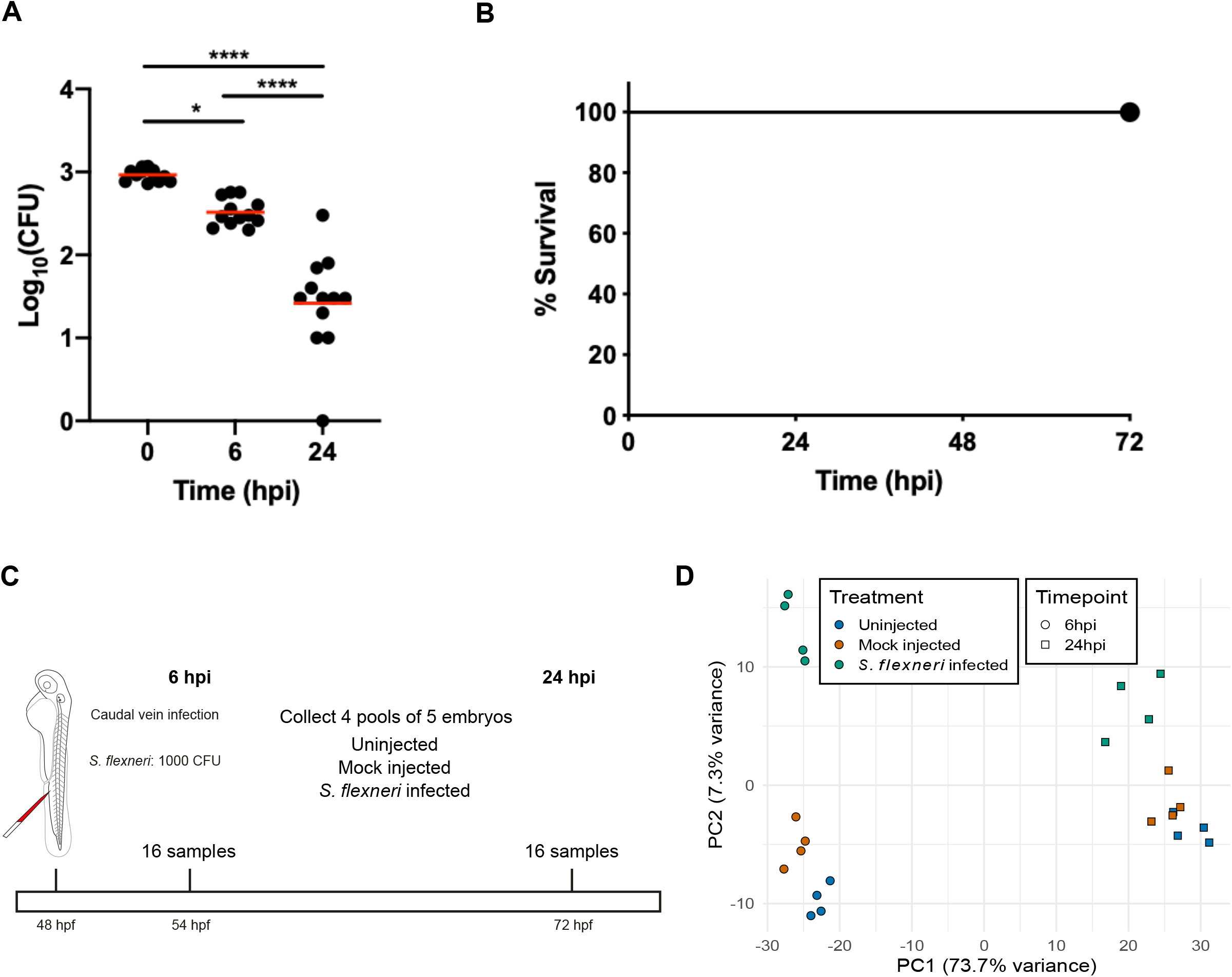
Experimental design of *Shigella*-zebrafish infection and transcriptomic data collection. **A-B**. Log10-transformed CFU counts (A) and survival curves (B) of larvae injected via the caudal vein with 1000 CFU of S. flexneri. Injections were performed in 2 day-post-fertilisation larvae. Statistics: one-way ANOVA with Tukey’s multiple comparisons test. *,p<0.05; ****,p<0.0001. A total of 36 larvae (12 per timepoint) were sacrificed to determine the bacterial load (A), while a total of 63 larvae were used for the survival analysis (B). **C**. Workflow of RNA sequencing experiment. 4 pools of 5 embryos injected with 1000 CFU of S. flexneri (or Mock-injected control) at 24 hours post-fertilisation were collected at 6 and 24 hours post-infection (hpi) for RNA sequencing. Uninfected embryos were also collected at the same timepoints. **D**. Principal component analysis (PCA). Regularized log-transformed counts for the 2500 most variable genes across the samples were used in PCA. The first two components are plotted. PC1 separates the samples by timepoint (circle=6 hpi, square=24 hpi) and PC2 reflects infection status (blue=uninfected, orange=mock infected, green=*S. flexneri* infected).

Principal component analysis (PCA) confirmed that biological replicates clustered according to their condition **(Fig. 1D)**. Stage explained the biggest principal component, accounting for 73.7% of the variance, and infection drove the second principal component, responsible for 7.3% of the variance. This shows that the largest source of variation in the data is between the two timepoints. The next largest is between uninfected and infected embryos at 6 hpi, suggesting the strongest response to infection is at 6 hpi. Overall, these data indicate a sharp distinction between the early (6 hpi) and late (24 hpi) responses to infection.

### Overview of RNA-seq results and gene ontology enrichment analysis

To identify the transcriptional signature of *S. flexneri* infection, we performed differential expression (DE) analysis between infected and mock-injected samples for *S. flexneri* infected larvae at 6 and 24 hpi. By comparing the two timepoints, we found a much stronger response at 6 hpi, where 1296 zebrafish genes were differentially expressed. By 24 hpi the expression level of these genes returned comparable to that of mock-injected larvae, and the overall number of DE genes decreased from 1296 to 111 genes (**Fig. 2A, B**). Additionally, the overlap between the two timepoints (i.e., genes that were consistently DE at both timepoints examined) was only 50 genes (**Fig. 2C**). Overall, these findings indicate that sublethal infections with *S. flexneri* are characterised by an acute and short-lived variation in gene expression, which subsides as bacterial load is cleared.

**Figure 2.**
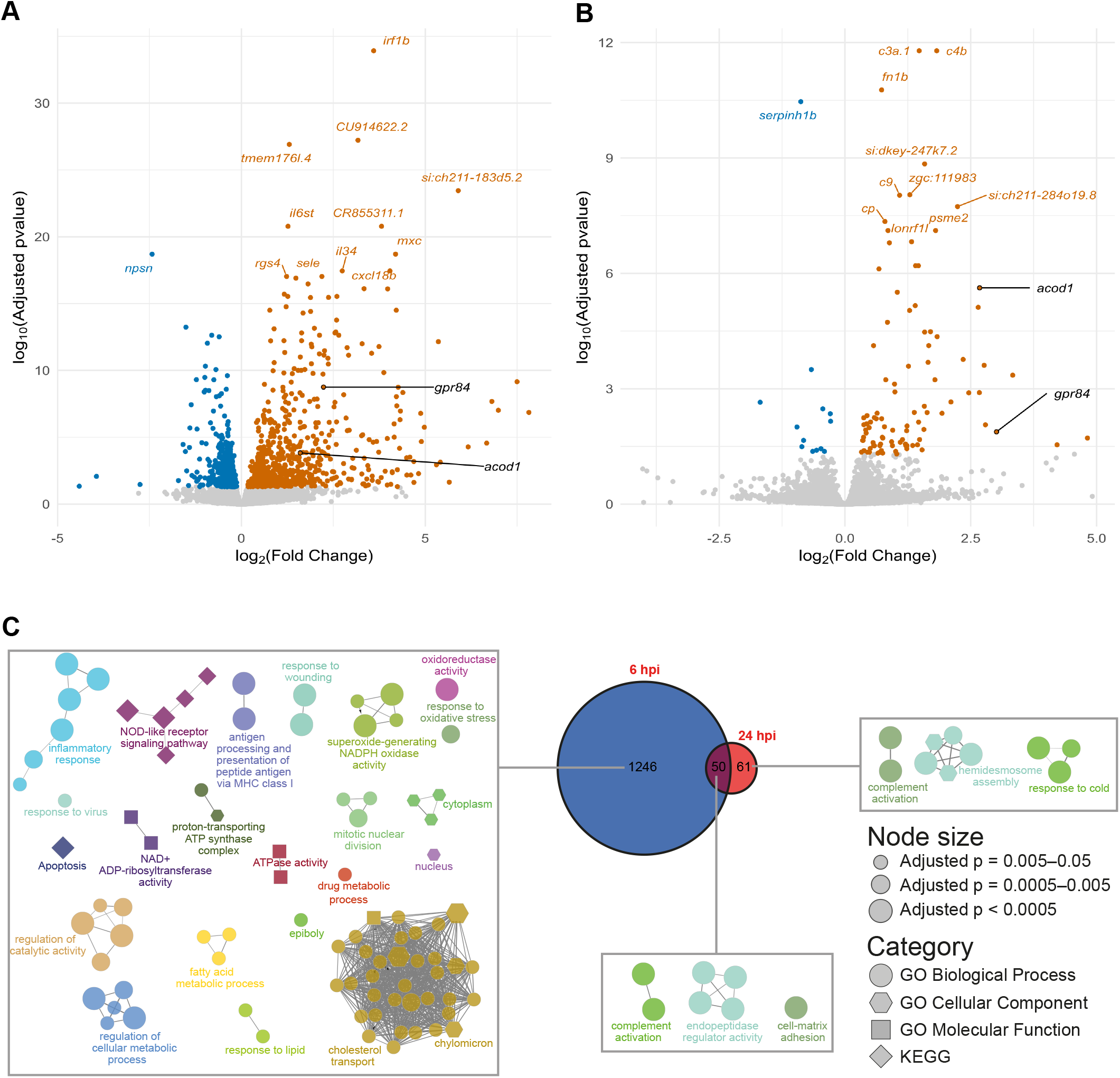
Analysis of differentially expressed genes in *S. flexneri* injected embryos. **A-B**. Volcano plots of differentially expressed genes between embryos infected with *S. flexneri* and mock-injected at 6 hpi (A) and 24 hpi (B). Each point represents a gene, - log_10_(Adjusted p-value) is plotted on the y-axis and log_2_(Fold Change) on the x-axis. Upregulated genes are coloured in orange and downregulated ones in blue. Genes with the highest –log_10_(Adjusted p-value) are labelled. *gpr84* and *acod1* are highlighted in black as these genes were further pursued for functional characterization for a role in susceptibility to infection **C**. Gene Ontology (GO) term enrichments. Network diagrams of GO term enrichments for genes differentially expressed at 6 hpi only (left), 24 hpi only (right) and both 6 and 24 hpi (middle). Each node in the diagrams represents an enriched GO term and terms are connected to terms that share annotated genes. This clusters the terms into process-related groups. The Venn diagram shows the numbers of differentially expressed genes at each timepoint and the overlap.

To gain insight into processes transcriptionally regulated in response to *S. flexneri* infection, we performed gene ontology (GO) enrichment analysis. For this, we analysed separately the DE gene lists produced at 6 and 24 hpi. We additionally performed GO analysis for the genes that represented the intersection of the two timepoints (**Fig. 2C**). At both 6 and 24 hpi the GO enrichment analysis was dominated by immune related GO terms, including “inflammatory response”, “NOD-like receptor signalling pathway” and “complement activation”.

We next analysed up- and down- regulated genes separately (**Fig. 3**). *S. flexneri* responses at 6 hpi presented significant upregulation of GO terms “inflammatory response”, “cytokine-mediated signalling pathway”, “innate immune response” and “response to bacterium” (**Fig. 3A**). In contrast, significantly downregulated GO terms did not include any strictly immune related terms, but included “cortical actin cytoskeleton organization” and “regulation of cytokinesis” (**Fig. 3A**), which are in line with the ability of *S. flexneri* to manipulate the host cytoskeleton (Mostowy and Shenoy, 2015).

**Figure 3.**
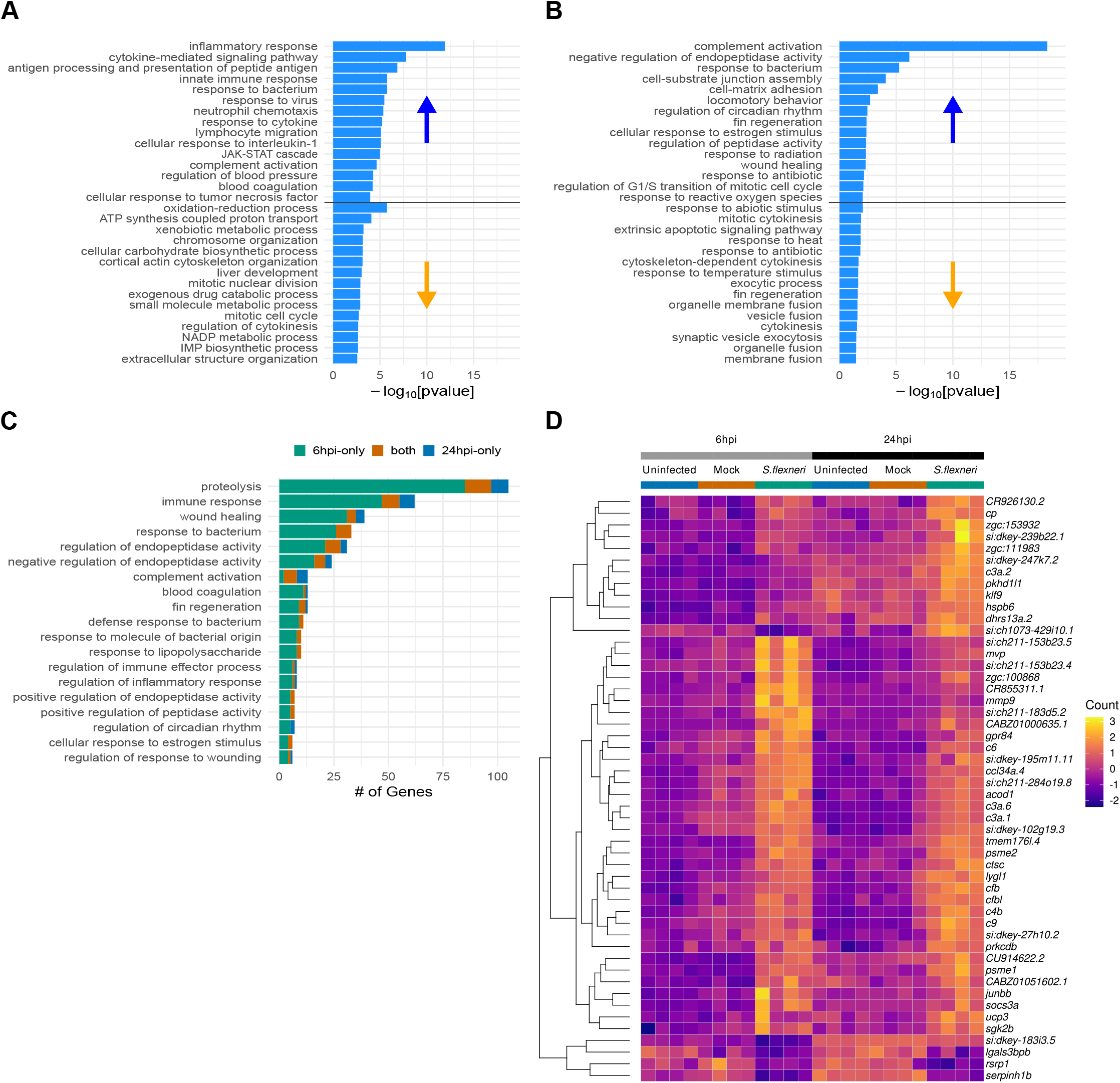
Gene ontology enrichment analysis of *S. flexneri* injected embryos. **A-B**. Histogram chart of the top 15 GO terms enriched by either up- or downregulated genes. A) 6 hpi. B) 24 hpi. Bars represent –log_10_[pvalue] for the enrichment. Each plot is divided into enrichments caused by upregulated genes (top half) and caused by downregulated genes (bottom half). **C**. Barchart of numbers of genes driving the enrichment of GO terms that are shared between the timepoints. Genes differentially expressed at 6 hpi only are shown in green, those at 24 hpi only in blue and those at both timepoints in orange. **D**. Heatmap of expression of the 50 genes that are differentially expressed at both 6 and 24 hpi in *S. flexneri* vs mock-infected embryos. The colour scale represents normalised counts calculated by DESeq2 that have been mean-centred and scaled by standard deviation for each gene across all the samples.

*S. flexneri* responses at 24 hpi presented significant upregulation of the GO terms “complement activation”, “response to bacterium” and “wound healing” (**Fig. 3B**). Significantly downregulated GO terms included “mitotic cytokinesis”/“cytokinesis” and “cytoskeleton dependent cytokinesis” (**Fig. 3B**). Despite overlap of enriched GO terms between 6 hpi and 24 hpi the genes driving the enrichments were not necessarily the same (**Fig. 3C**).

Only 50 genes were commonly differentially expressed irrespective of time point (**Fig. 3D** heatmap). These genes included several complement factors (*cfb, c4b, c3a*.*6, c3a*.*1*), *mmp9* (matrix metalloproteinase 9), *lygl1* (lysozyme g-like 1), *psme2* (proteasome activator complex subunit 2), *ccl34a*.*4* (C-C motif chemokine ligand 34a.4), *acod1* (aconitate decarboxylase 1) and *gpr84* (G-protein coupled receptor 84).

Together, whole animal RNA-seq profiling identified a novel set of markers of *S. flexneri* infection and zebrafish host defence (**Table S1**).

### Discovery of host factors controlling *S. flexneri* infection

To test the roles of select DE genes we took advantage of previously published mutants and ENU-induced mutations from the Zebrafish Mutation Project (ZMP) (Kettleborough, Busch-Nentwich, Harvey, Dooley, De Bruijn, *et al*., 2013). We also generated new mutants using CRISPR/Cas9 mutagenesis (Brocal *et al*., 2016).

We first established a method to screen for susceptibility phenotypes, using the previously published *irf8* mutant line (Shiau *et al*., 2015). *Irf8* mutants have significantly reduced macrophage numbers, and we previously showed that *irf8* depletion (by morpholino oligonucleotide) and macrophage ablation (by metronidazole treatment in a transgenic line expressing the nitro-reductase enzyme in macrophages) increase susceptibility to *S. flexneri* (Mazon-Moya *et al*., 2017; Torraca *et al*., 2019). In line with these observations, homozygous *irf8* mutants are significantly more susceptible to *S. flexneri* than their wild-type siblings (**Fig. 4A**). From this, we concluded that our experimental approach from RNA-seq profiling to zebrafish survival is highly suited to discover genes controlling susceptibility to *S. flexneri* infection.

**Figure 4.**
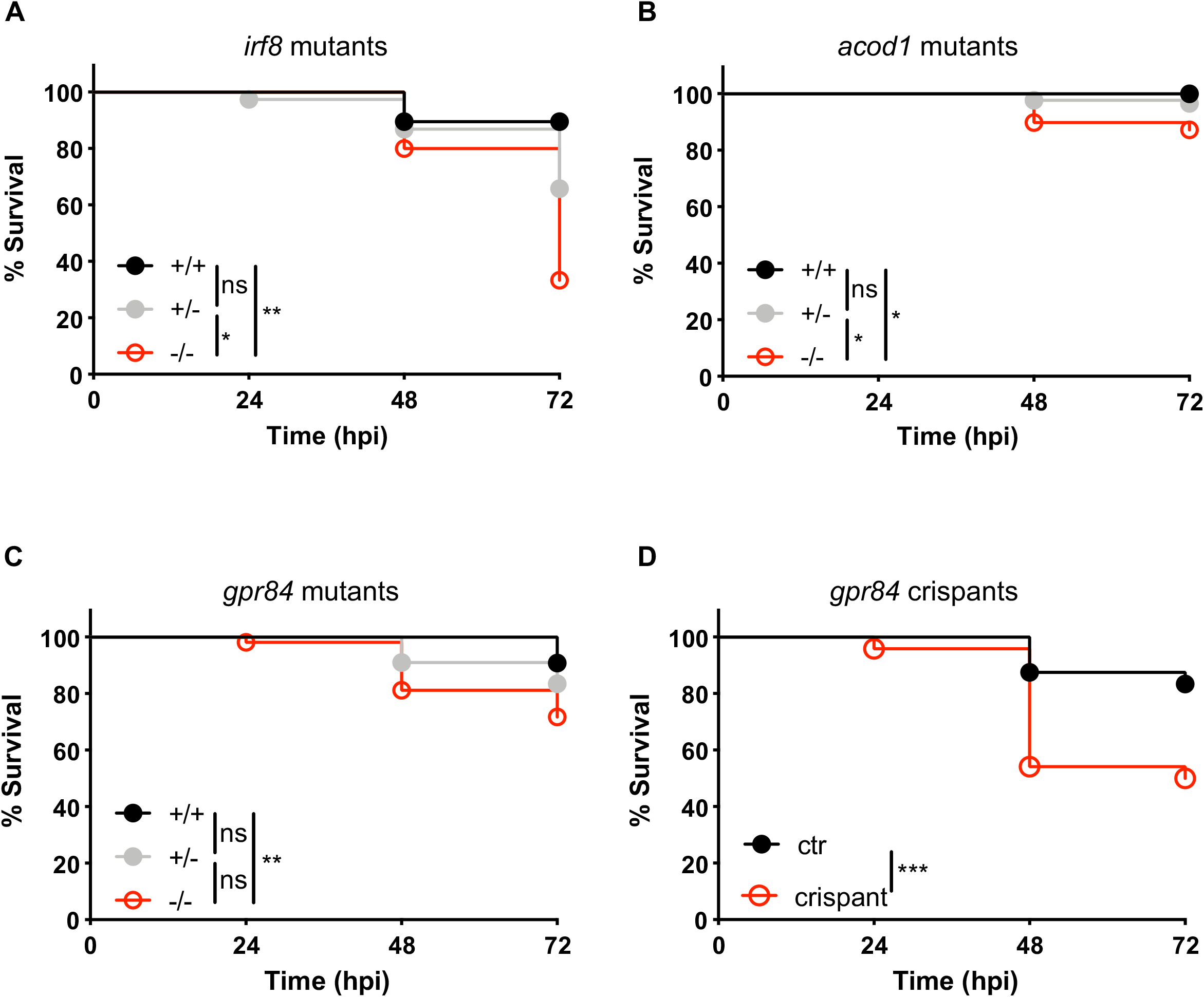
Functional analysis of zebrafish mutants in susceptibility to infection. Survival curves of larvae derived from incrosses of *irf8* (A), *acod1* (B), or *gpr84* (C) heterozygous carriers of null mutations and survival curves of *gpr84* crispant larvae or controls (D). Injections were performed in 2 day-post-fertilisation larvae via the hindbrain ventricle with 5.000-10.000 CFU of *S. flexneri*. Statistics: Log-rank (Mantel-Cox) test. For A-C, all larvae were individually genotyped post-mortem or at the end of the experiment. For D, a few randomised larvae were individually genotyped to confirm efficient CRISPR targeting *,p<0.05; **,p<0.01; ***,p<0.001. A total of 72 (A), 172 (B), 262 (C) or 96 (D, 48 crispants and 48 controls) larvae were used for the survival analyses.

Next, we explored the role of *acod1 (*also known as *irg1, immunoresponsive gene 1)* and *gpr84* in the context of *S. flexneri* infection, since these two factors are among the few genes that are consistently induced by *S. flexneri* at both time points. Strikingly, homozygous mutants for the *acod1*^*sa31153*^ or the *gpr84*^*sa10052*^ allele are significantly more susceptible to *S. flexneri* infection than their wild-type siblings (**Fig. 4B-C**). The role of *gpr84* in controlling *S. flexneri* infection was confirmed by F0 CRISPR/Cas9 mutagenesis, as *gpr84* crispants phenocopied *gpr84* homozygous mutants (**Fig. 4C-D**).

## DISCUSSION

The zebrafish larva is a powerful non-mammalian vertebrate model, that has been particularly instrumental to study the innate immune response to bacterial infection (Torraca *et al*., 2018; Gomes and Mostowy, 2020; Stream and Madigan, 2022). We previously used *S. flexneri* infection of zebrafish to study bacterial autophagy (Mostowy et al, 2013), predator-prey interactions (Willis et al, 2016), septin-mediated immunity (Mazon Moya 2017), innate immune training (Willis et al 2018), *Shigella* evolution (Torraca *et al*., 2019) and septin-mediated apoptosis (Van Ngo et al 2022). Here, using RNA-seq we transcriptionally capture the host response to *S. flexneri* infection over time. We hope this work will serve as reference for future *in vivo* studies of *S. flexneri* infection. Moreover, it is envisioned that discoveries from genome-wide host RNA signatures of infectious diseases can be used for clinical translation.

Our transcriptional profiles reveal that *acod1* and *gpr84* are upregulated at both 6 and 24 hpi, suggesting a lasting role during *S. flexneri* infection. Consistent with this, we show that zebrafish mutants in these genes are significantly more susceptible to infection. In the case of *acod1*, similar results were previously published using *Salmonella* Typhimurium (a primary enteric pathogen infecting humans and other animals) (Hall *et al*., 2013). While the role of complement factors, chemokines, hydrolytic enzymes and aconitate dehydrogenase have been associated with a variety of infectious and inflammatory processes (Murdoch and Finn, 2000; Elkington, O’Kane and Friedland, 2005; Dunkelberger and Song, 2009; Ragland and Criss, 2017; Sommer, Torraca and Meijer, 2020; Wu, Kang and Tang, 2022), *gpr84* association to infection was mostly unknown. Zebrafish Gpr84 protein sequence shares 51% homology with the human GPR84 protein (**Suppl. Fig. 1**), and *in vitro* studies showed that GPR84 is upregulated in macrophages and neutrophils after lipopolysaccharide (LPS) stimulation (Lattin *et al*., 2008). Considering this, it is next of great interest to understand the interplay between *gpr84* and LPS in controlling *S. flexneri* infection.

## ACKNOWLEDGEMENTS

We thank Gordon Dougan for encouraging this work. We thank Antonio Pagán for help with the crispant methodology and for suggestions regarding the survival experiments. V.T. was supported by the European Union’s Horizon 2020 research and innovation program under the Marie Skłodowska-Curie (H2020-MSCA-IF-2015 – 700088) and an LSHTM/Wellcome Institutional Strategic Support Fund (ISSF) Fellowship (204928/Z/16/Z). Research in the Mostowy laboratory was supported by a Wellcome Trust Research Career Development Fellowship (WT097411MA) and the Lister Institute of Preventive Medicine, and is supported by a Wellcome Trust Senior Research Fellowship (206444/Z/17/Z) and a European Research Council Consolidator Grant (772853 - ENTRAPMENT).

## MATERIAL AND METHODS

### Ethics statements

Animal experiments were performed according to the Animals (Scientific Procedures) Act 1986 and approved by the Home Office (Project licenses: PPL P84A89400 and P4E664E3C). All experiments were conducted up to 5 days post fertilisation.

### Zebrafish

Zebrafish lines used here were the wildtype (WT) AB strain. The Sanger mutants were made originally in the wild-type TL background. *acod1*^*sa31153*^ was maintained in a mixed AB/TL background, while *gpr84*^*sa10052*^ (Kettleborough, Busch-Nentwich, Harvey, Dooley, de Bruijn, *et al*., 2013) was maintained in the TL background. Unless specified otherwise, eggs, embryos and larvae were reared at 28.5°C in Petri dishes containing embryo medium, consisting of 0.5x E2 water supplemented with 0.3 μg/ml methylene blue (Sigma-Aldrich, St. Louis, Missouri). For injections, anaesthesia was obtained with buffered 200 μg/ml tricaine (Sigma-Aldrich) in embryo medium. Protocols are in compliance with standard procedures as reported at zfin.org. For *acod1* and *gpr84* mutants, genotyping was performed using the KASP assay (LGC Biosearch technologies). For *irf8* mutants and *gpr84* crispants, genotyping was performed using a high-resolution melting curve analysis (Garritano *et al*., 2009; Pagan *et al*., 2015). Primers used for the assays are reported in **Table 1**.

### F0 CRISPR mutagenesis

*The acod1*^*sa31153*^ allele was created using CRISPR-Cas9 mutagenesis as described (Brocal et al., 2016). The CRISPR guide RNA was targeted to chr9:21836233-21836255 (GRCz11). The guide RNA sequence is TCCAGGCCAGAGGGTTTAAC<u>AGG</u> (PAM underlined). *sa31153* is a 35 bp deletion from 21836248–21836282 (deleted sequence is TTAACAGGCAGTGTTTCAGCCAGTGCCAGGAGAGC). This leads to a frameshift that is predicted to result in a stop codon after 50 amino acids. Guide RNAs for injection were produced *in vitro* following the method of Hwang et al. (Hwang *et al*., 2013). Briefly, overlapping oligonucleotides for the target site (IDT) were annealed and cloned into the pDR274 vector (Addgene: 42250) linearised with BsaI (New England Biolabs). Sequence-confirmed vectors were then linearized with DraIII (New England Biolabs), and sgRNA transcripts were generated using the MEGAshortscript T7 Kit (ThermoFisher Scientific). sgRNAs were then DNase treated and precipitated with ammonium acetate/ethanol. Cas9 mRNA was transcribed from linearised pCS2-nls-zCas9-nls plasmid (Addgene: 47929) (Jao, Wente and Chen, 2013) using mMessage Machine SP6 kit (ThermoFisher Scientific), DNase treated and purified by phenol-chloroform extraction and EtOH precipitation. RNA concentration was quantified using Qubit spectrophotometer. Approximately 1 nl total volume of 10 ng/ul (sgRNAs) and 200 ng/ul (Cas9 mRNA) was injected into the cell of one-cell stage embryos. Embryos were raised and screened as detailed in Brocal et al. (Brocal *et al*., 2016) to isolate carriers.

### Bacterial infections

GFP fluorescent or non-fluorescent *S. flexneri* M90T was used for all infections. Bacterial preparation and infection were described previously (Torraca *et al*., 2019). For all experiments, 1–2 nl of bacterial suspension (bacterial load as indicated in the individual experiments) or control solution were microinjected in the caudal vein or the hindbrain ventricle (HBV) of 2 days post-fertilisation (dpf) zebrafish larvae, as specified in the individual figure legend. Bacterial enumeration was performed *a posteriori* by mechanical disruption of infected larvae in 0.4% Triton X-100 (Sigma-Aldrich) and plating of serial dilutions onto Congo red-TSA plates. No significant effect in survival was observed between mutants/crispants and their wild-type siblings when embryos were challenged with mock injections.

### RNA sequencing

RNA was extracted from larvae as described previously (Wali *et al*., 2022). Briefly, RNA was extracted from individual embryos by mechanical lysis in RLT buffer (Qiagen) containing 1 μl of 14.3 M β-mercaptoethanol (Sigma). The lysate was combined with 1.8 volumes of Agencourt RNAClean XP (Beckman Coulter) beads and allowed to bind for 10 min. The plate was applied to a plate magnet (Invitrogen) until the solution cleared and the supernatant was removed without disturbing the beads. This was followed by washing the beads three times with 70% ethanol. After the last wash, the pellet was allowed to air-dry for 10 min and then resuspended in 50 μl of RNAse-free water. RNA was eluted from the beads by applying the plate to the magnetic rack. Samples were DNase-I treated to remove genomic DNA. RNA was quantified using Quant-IT RNA assay (Invitrogen). ERCC spike mix 2 (Ambion) was added to the RNA. Stranded RNA-seq libraries were constructed using the Illumina TruSeq Stranded RNA protocol after treatment with Ribozero. Libraries were pooled and sequenced on Illumina HiSeq 2500 in 75 bp paired-end mode (median 7.3 million reads per sample). Sequence data were deposited in ENA under accession ERP012128. The data were assessed for technical quality (GC-content, insert size, proper pairs, etc.) using FASTQC (https://www.bioinformatics.babraham.ac.uk/projects/fastqc/). Reads for each sample were mapped to the GRCz11 zebrafish genome assembly and counted against the Ensembl version99 annotation using STAR (Dobin *et al*., 2013). Differential expression analysis was done in R (R Core Team, 2019) with DESeq2 (Love, Huber and Anders, 2014) using a cut-off for adjusted p-values of 0.05.

## Data Availability

Sequence data are available in ENA under accession ERP012128. The output from DESeq2 for each timepoint is in **Table S1** (doi.org/10.6084/m9.figshare.20768851).

## FIGURE LEGENDS

**Supplementary Figure 1. *In silico* analysis of zebrafish Grp84 and comparison to human and murine Gpr84**.

**A**. Phylogeny tree for Gpr84, showing clustering of *Danio rerio* (dre) *Mus musculus* (mmu) and *Homo sapiens* (hsa) protein sequences. Protein sequence for human Adra1 (Alpha-1A Adrenergic Receptor) was the closest protein sequence to human Gpr84 within the human genome and was used to root the tree. Phylogeny tree was obtained by ClustalO, using protein sequences for the canonical protein isoform or the predicted principal isoform. The tree was edited and rooted using figtree.

**B**. Zebrafish gpr84 is predicted to have a 7-loop transmembrane architecture by TMHMM - 2.0, which represents the typical architecture of a G-protein coupled receptor.

**C**. Sequence alignment of *Danio rerio* (dre) *Mus musculus* (mmu) and *Homo sapiens* (hsa) Gpr84 protein sequences. Alignment was obtained by ClustalO (https://www.ebi.ac.uk/Tools/msa/clustalo/) and visualised using MView (https://www.ebi.ac.uk/Tools/msa/mview/).

**Table S1. List of differentially expressed genes in *S. flexneri* injected embryos** Differential expression analysis was done in R with DESeq2 using a cut-off for adjusted p-values of 0.05.

